# Foundry-fabricated dual-color nanophotonic neural probes for photostimulation and electrophysiological recording

**DOI:** 10.1101/2024.09.25.614961

**Authors:** David A. Roszko, Fu-Der Chen, John Straguzzi, Hannes Wahn, Alec Xu, Blaine McLaughlin, Xinxin Yin, Hongyao Chua, Xianshu Luo, Guo-Qiang Lo, Joshua H. Siegle, Joyce K. S. Poon, Wesley D. Sacher

## Abstract

**Significance:** Compact tools capable of delivering multicolor optogenetic stimulation to deep tissue targets with sufficient span, spatiotemporal resolution, and optical power remain challenging to realize. Here, we demonstrate foundry-fabricated nanophotonic neural probes for blue and red photostimulation and electrophysiological recording, which use a combination of spatial multiplexing and on-shank wavelength-demultiplexing to increase the number of on-shank emitters.

**Aim:** We demonstrate Si photonic neural probes with 26 photonic channels and 26 recording sites, which were fabricated on 200-mm diameter wafers at a commercial Si photonics foundry. Each photonic channel consists of an on-shank demultiplexer and separate grating coupler emitters for blue and red light, for a total of 52 emitters.

**Approach:** We evaluate neural probe functionality through bench measurements and *in vivo* experiments by photostimulating through 16 of the available 26 emitter pairs.

**Results:** We report neural probe electrode impedances, optical transmission, and beam profiles. We validated a packaged neural probe in optogenetic experiments with mice sensitive to blue or red photostimulation.

**Conclusions:** Our foundry-fabricated nanophotonic neural probe demonstrates dense dual-color emitter integration on a single shank for targeted photostimulation. Given its two emission wavelengths, high emitter density, and long site span, this probe will facilitate experiments involving bidirectional circuit manipulations across both shallow and deep structures simultaneously.

## 1 Introduction

Optogenetics has enabled neuroscientists to selectively excite, inhibit, and modulate neural activity in a cell-type specific, wavelength-selective manner. Today, a wide range of optogenetic actuators are available, with peak activation wavelengths across the visible spectrum.^1–8^ In combination, these optogenetic actuators can enable diverse experimental designs, as opsins with distinct activation spectra (i.e., blue- and red-light activated opsins) can be used for independent excitation and inhibition of different cell types within the same region^9^ or excitation and inhibition of the same cell type.^8^

To facilitate optogenetic experiments, it is crucial to develop tools that can deliver multicolor photostimulation with high spatiotemporal resolution while simultaneously recording evoked neural activity. For delivering light into superficial brain regions (depth: *≤*1 mm), methods based on two-photon microscopy have been widely used.^10, 11^ However, to interrogate deeper neural tissue (depth: *>*1 mm), researchers have developed implantable tools based on integrated optoelectronic and photonic technologies. These approaches are based on microchips consisting of (1) long (3-10 mm), narrow, implantable shanks with integrated optoelectronic or photonic emitters for photo-stimulation and (2) a larger base region with circuitry and electrical/optical interfaces. Each of these technologies exhibit notable trade-offs in terms of average emitter power, emitter numbers, and on-shank emitter densities.

Optoelectronic neural probes with high-density on-shank integrated micro light-emitting diodes (*µ*LEDs) or organic light-emitting diodes (OLEDs) have been developed for dual-color photostimulation and neural recording.^12–14^ Recent reports have demonstrated high emitter counts (16 red and blue *µ*LEDs per shank in Ref. 12, 256 orange or blue OLEDs per shank in Ref. 13). However, these designs either have large shank dimensions (200 *µ*m wide in Ref. 12), which may cause excess tissue damage, or low output powers (40-100 nW in Ref. 13), which necessitate using multiple emitters for photostimulation with most opsins - effectively reducing the spatial resolution of the device. Additionally, designs typically leverage custom fabrication steps beyond those used for CMOS electronics, posing challenges for scaling manufacturing volumes.

Photonic neural probes guide light from external laser sources into the brain via integrated optical waveguides and diffractive (grating coupler) emitters. By delocalizing the light source from the shank and surrounding brain tissue, photonic probes offer the opportunity for high output powers beyond those of optoelectronic probes, which are limited by on-shank emitter heating. This approach also leverages the increasing maturity of silicon (Si) integrated photonics technology, charting a path toward high emitter counts on narrow shanks using dense arrays of nanophotonic waveguides in addition to scalable manufacturing volumes via commercial Si photonics foundries. Recent reports of photonic probes most often rely on silicon nitride (SiN) waveguides,^15–19^ which are commonly used in standard Si photonics^20^ and can be engineered for broadband transparency extending to the visible spectrum.^19, 21^ This broadband transparency is of particular utility for multiplexing multiple photostimulation wavelengths onto each waveguide.

Currently, photonic neural probe technology is in its early development stages, with low channel counts relative to their optoelectronic probe counterparts and few reports of dual color photostimulation functionalities. An early example of a dual-color photonic probe used silicon oxynitride waveguides to guide 405 and 635 nm light, but the large waveguide cross sectional areas (7×30 *µ*m^2^) limited the design to one emitter per shank.^22^ More recent demonstrations with sub-micron SiN waveguides have achieved higher channel counts. In Ref. 17, 3 red and 3 blue emitters were integrated onto a single shank and coupled to a single bus waveguide via ring resonator wavelength demultiplexer devices. Tuning of the input wavelength enabled emitter selection (wavelength-division multiplexing). In this approach, scaling of the number of emitters (and ring resonators) is expected to be limited by the inherent fabrication variation sensitivity of ring resonator devices.^23^ In Ref. 19, 14 red and 14 blue emitters were integrated onto the shank of a CMOS-based neural probe, with each emitter coupled to a separate waveguide. Emitters were selected by an on-chip photonic switching circuit integrated onto the base of the probe (spatial multiplexing). With spatial multiplexing alone, emitter densities are limited by parasitic optical coupling/crosstalk between adjacent waveguides.

Here, we present dual-color photonic neural probes with combined spatial multiplexing and wavelength-division multiplexing for increasing the number of addressable emitters on a single shank. Our design, which builds on previous single-wavelength integrated probe designs from our group,^18^ is manufactured using wafer-scale processes at a commercial Si photonics foundry (Advanced Micro Foundry, Singapore), paving the way for mass production and broad dissemination within the neuroscience community. Each neural probe has 52 emitters on a single shank (26 blue and 26 red), with emitter addressing via both spatial multiplexing (with external laser scanning optics addressing a multicore fiber coupled to the probe) and wavelength multiplexing (with red and blue input lasers). An array of spatially-addressed, fiber-to-chip edge couplers and waveguides -each coupled to a core of the multicore fiber - route multiplexed blue (473 nm) and red (638 nm) light to on-shank evanescent directional-coupler wavelength demultiplexers, which direct each color to separate (red- or blue-designed) grating coupler emitters. Additionally, our design contains 26 recording electrodes for electrophysiological recording. We demonstrate the probe experimentally through *in vivo* experiments in optogenetic mice sensitive to either blue or red photostimulation. A conceptual overview is shown in Fig. 1. Compared to prior reports of dual-color photonic neural probes,^17, 19, 22^ the neural probes demonstrated herein achieve a record number of emitters per shank. Furthermore, this work highlights the potential of combined spatial- and wavelength-multiplexing for scaling to high-channel count, high-density dual-color photonic neural probes.

**Fig 1.**
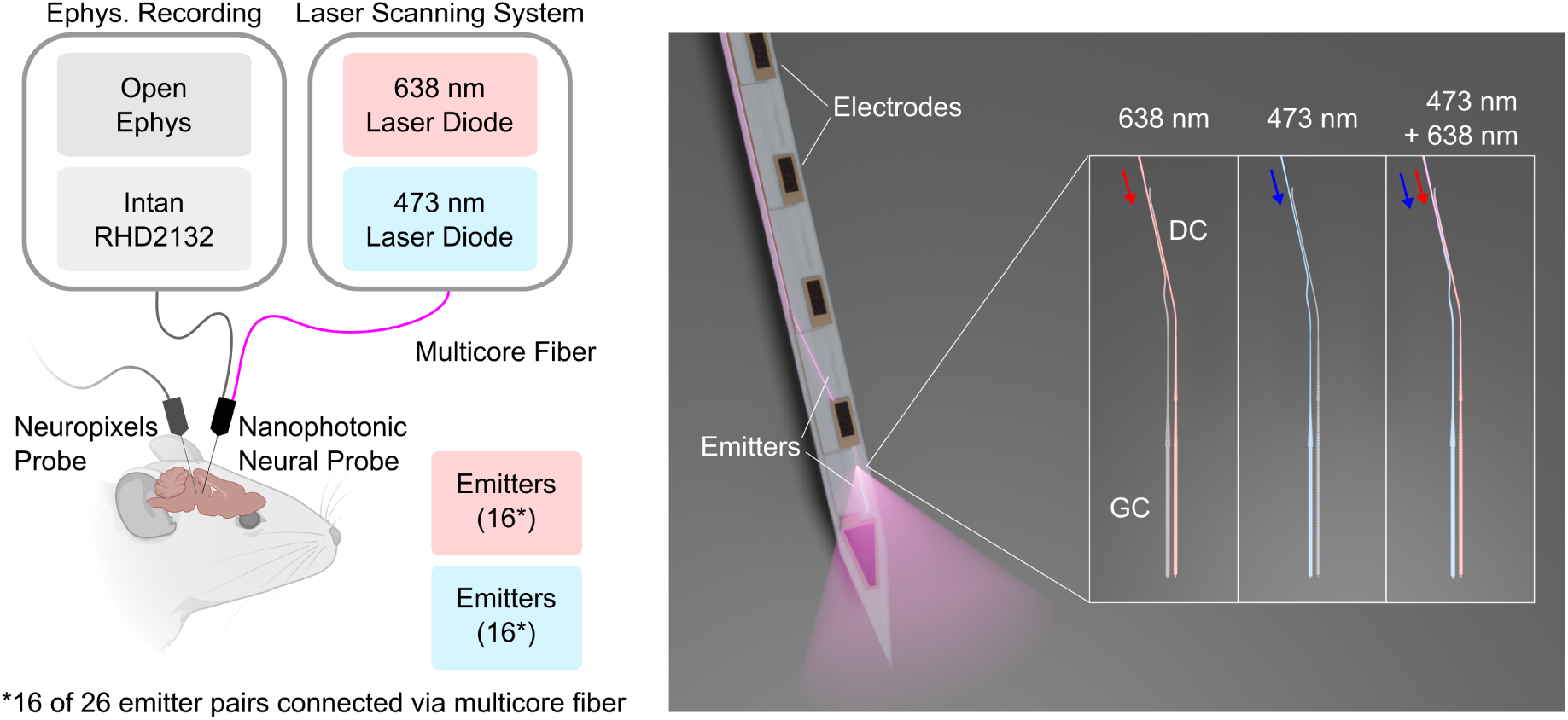
Left: Conceptual figure illustrating simultaneous recordings with a dual-color nanophotonic neural probe and a Neuropixels probe.^24^ Figure created with BioRender.com. Right: Graphical rendering of a dual-color nanophotonic probe. Each routing waveguide connects to a directional coupler (DC) which demultiplexes red (638 nm) and blue (473 nm) light to separate grating coupler (GC) emitters.

## 2 Results

### 2.1 Neural Probe Design and Characterization

Neural probes were designed and simulated using Lumerical finite-difference time-domain (FDTD) optical simulations (Ansys, Inc., Canonsburg, PA, USA). Final designs were fabricated by Advanced Micro Foundry (AMF) in Singapore using a custom visible-light SiN photonic process.^25^ Wafers were received from AMF (Fig. 2a) and chips were removed for testing and further processing. The wafer cross-section consisted of two SiN layers (SiN1 and SiN2) with SiO_2_ cladding for photonic routing and 3 aluminum (Al) metal layers for recording electrophysiological activity (Fig. 2b). Recording electrodes were implemented by coating the top aluminum layer with TiN. For the neural probes in this study, we aimed to design a probe with sufficiently long span and emitter density to interrogate superficial and deep neural targets with blue (473 nm) and red (638 nm) light. The neural probes, shown in Fig. 2c, were designed with a 6.09 mm long and 70 *µ*m wide shank. 25 TiN recording electrodes (46.5 um x 14 um), with 1 additional TiN reference electrode at the shank tip, were distributed linearly along the shank with a 188 *µ*m pitch and span of 4.80 mm (Fig. 2d). An array of 26 fiber-to-chip bilayer edge couplers^26^ were fabricated at the base of the device for coupling light from external lasers using a multicore fiber. To achieve high-density emitters on the shank, our design implemented a wavelength multiplexing scheme whereby each edge coupler interfaced with a nanophotonic waveguide capable of routing both 473 nm and 638 nm light down the shank of the probe. To reduce waveguide crosstalk, neighbouring single-mode waveguides were adiabatically transitioned to dissimilar (detuned) multimode widths of 600 nm and 700 nm.^27^ On the shank, each routing waveguide terminates with an evanescent directional coupler wavelength demultiplexer designed to filter 473 nm and 638 nm light to wavelength-specific bilayer grating coupler emitters, consisting of a fully-etched grating in SiN2 and a corrugated grating^28^ in SiN1 (Fig. S1). Overall, each neural probe has 26 routing waveguides which route red and blue light to 26 pairs of grating coupler emitters, for a total of 52 grating coupler emitters. The pitch and span of the grating coupler emitter pairs were 188 *µ*m and 4.82 mm, respectively.

**Fig 2.**
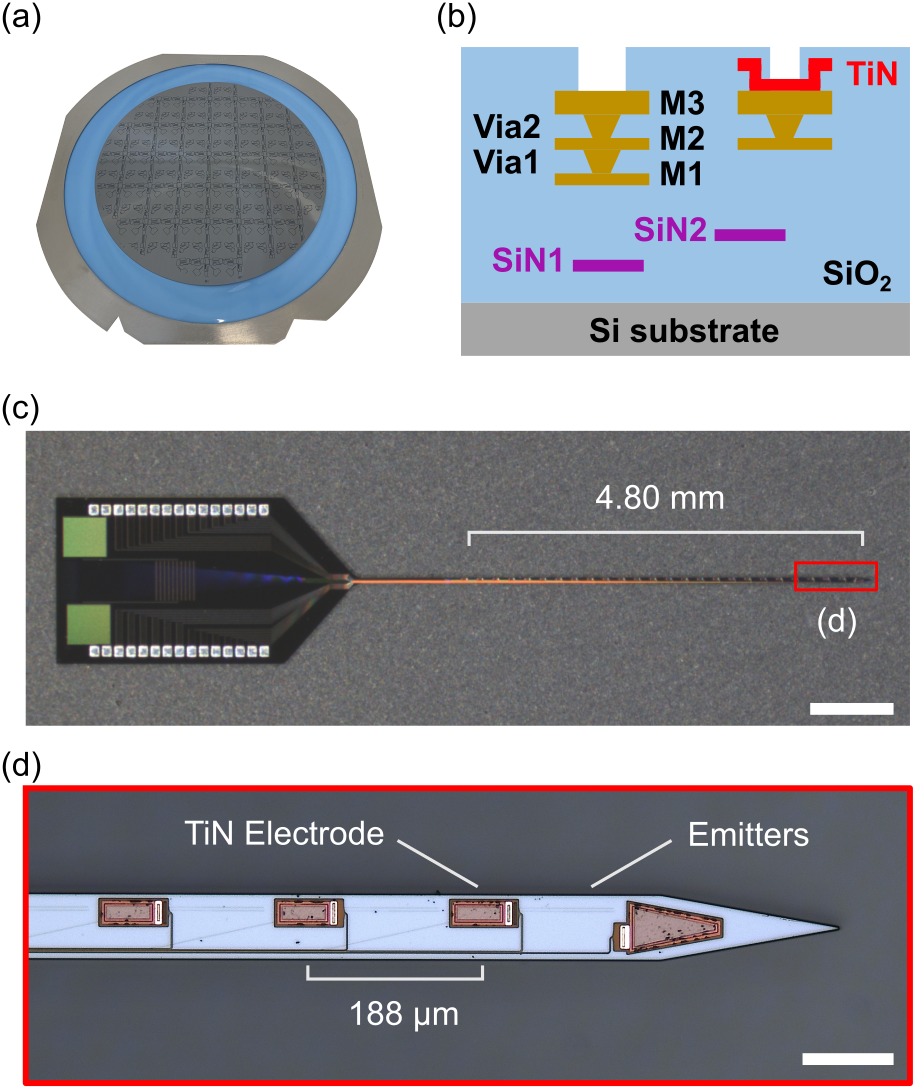
(a) Foundry-fabricated, 200-mm diameter, Si photonic neural probe wafer. (b) Cross-sectional schematic of the neural probes. (c) Micrograph of a fabricated dual-color nanophotonic neural probe. Scale bar: 1 mm (d) Enhanced micrograph showing TiN electrodes and emitters on the neural probe shank. Scale bar: 100 *µ*m

The optical transmission of neural probe samples prior to packaging (n=78 emitter pairs from 3 probes) and after packaging (n=64 emitter pairs from 4 probes) were measured and shown in Fig. 3a (left). We define the probe transmission as the ratio of output power from a grating coupler emitter to the input power incident at the edge coupler facet. Prior to packaging, the average probe transmission was measured as -12.6 *±* 1.2 dB (mean *±* SD) and -20.4 *±* 1.0 dB for 473 nm and 638 nm transverse electric (TE)-polarized light, and -15.8 *±* 1.2 dB and -25.6 *±* 0.6 dB for 473 nm and 638 nm transverse magnetic (TM)-polarized light, respectively. After packaging, transmission was measured using depolarized light (equal parts TE and TM on-chip) to mimic the conditions of the final experiments (see Methods 4.7), wherein temperature- and stress-induced polarization fluctuations of the multicore fiber coupled to the probe were avoided via depolarized input light. The average transmission for the packaged probes was measured as -27.1 *±* 4.7 dB and -29.5 *±* 4.3 dB for 473 nm and 638 nm light, respectively. A micrograph showing 473 nm and 638 nm light emitting from the deepest pair of grating coupler emitters can be seen in Fig. 3a (right).

**Fig 3.**
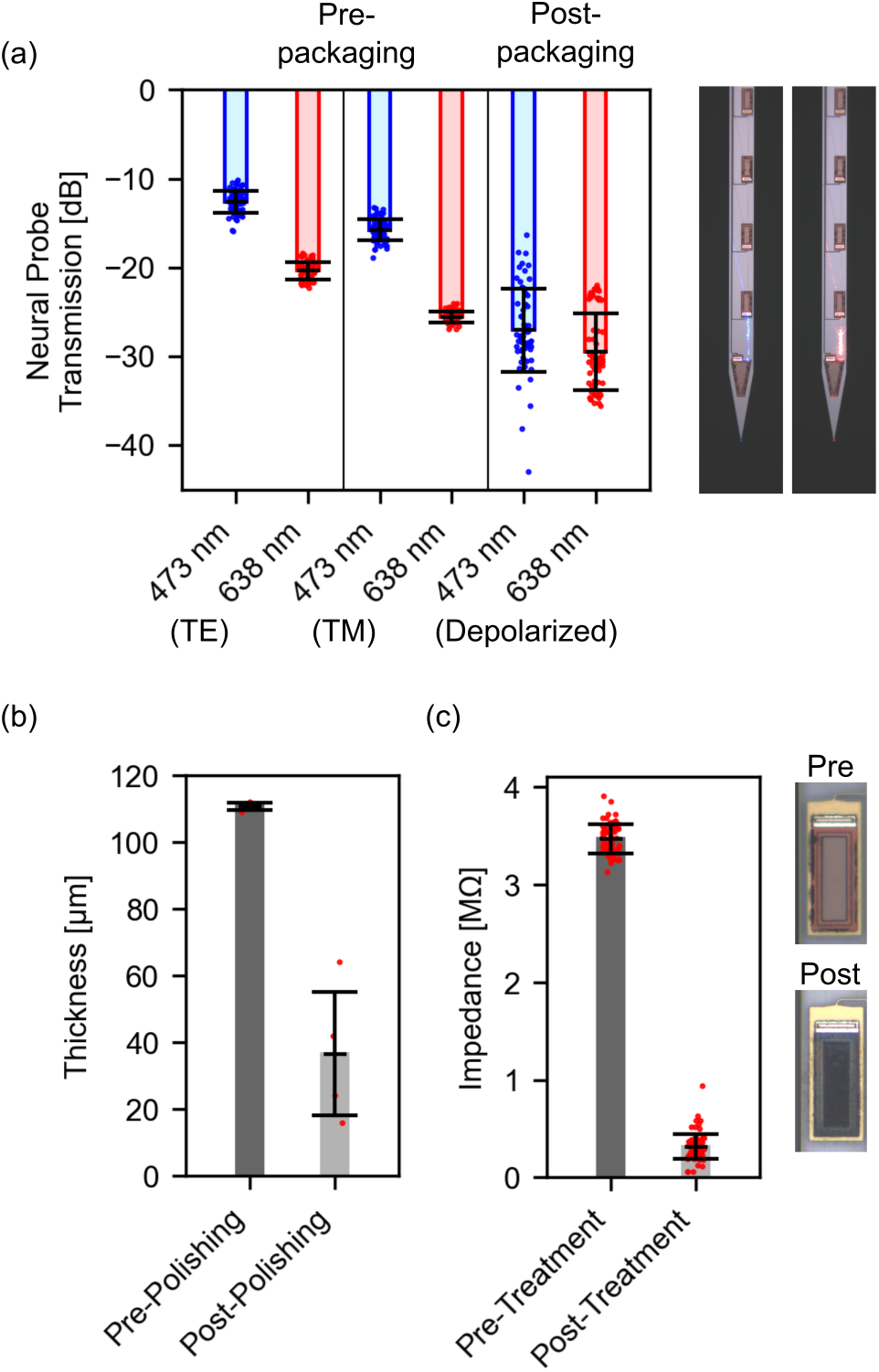
(a) Neural probe transmission measurements for blue (473 nm) and red (638 nm) light before and after packaging. (Pre-packaging TE and TM measurements: n=78 emitter pairs from 3 probes; post-packaging depolarized measurements: n=64 emitter pairs from 4 probes). Micrographs show blue and red light emitting from the deepest emitter pair on the neural probe. (b) Neural probe shank thickness before and after mechanical polishing (n=4 probes). (c) Electrode impedance magnitudes before and after laser-induced impedance reduction. Micrographs show sample electrodes before and after laser surface roughening (pre-treatment: n=75 electrodes from 3 probes; post-treatment: n=90 electrodes from 4 probes).

Neural probes were mechanically polished to reduce the overall device thickness in an effort to reduce tissue insertion damage during *in vivo* experiments. The thickness of select neural probe samples before and after mechanical polishing is shown in Fig. 3b. Prior to mechanical polishing the thickness of the extracted neural probe chips was 111 *±* 1 *µ*m (n=4). After mechanical polishing, the average thickness of the neural probes was reduced to 37 *±* 18 *µ*m (n=4).

The electrochemical impedance of the recording electrodes was reduced to improve electrophysiological signal recording during *in vivo* experiments. Electrochemical impedance reduction was achieved using selective femtosecond laser scanning with a 2-photon microscope to roughen the electrode surfaces, as described in Methods 4.2 and our previous work in Ref. 18. The results of the impedance reduction are shown in Fig. 3c (left). The impedance magnitude of the untreated TiN electrodes was measured as 3.47 *±* 0.15 MΩ (n=75 electrodes from 3 probes). After impedance reduction, the impedance magnitude of the TiN electrodes was measured as 315 *±* 125 kΩ (n=90 electrodes from 4 probes). Representative images showing the TiN electrodes before and after laser impedance reduction are shown in Fig. 3c (right).

### 2.2 On-shank directional-coupler wavelength-demultiplexer performance and grating coupler emitter beam profiles

Directional coupler demultiplexer (demux) test structures (n=5) were measured to evaluate how well 473 nm and 638 nm light were coupled to the appropriately designed grating coupler emitters (Fig. 4). Demux test structures were measured using both TE- and TM-polarized light. For each quadrant of Fig. 4, port 1 represents the output port designed to pass 473 nm light, and port 2 represents the output port designed to pass 638 nm light. For TE-polarized light (Figs. 4a and 4b), the transmission of port 1 was measured as -0.6 *±* 0.5 dB and -26.1 *±* 2.7 dB for 473 nm and 638 nm light, respectively, while the transmission of port 2 was measured as -15.8 *±* 0.7 dB and -3.3 *±* 1.5 dB for 473 nm and 638 nm light, respectively. On average, for TE-polarized light, the extinction ratio between the ports at 473 nm was 15.2 dB, while the extinction ratio at 638 nm was 22.8 dB. Similarly, for TM-polarized light (Figs. 4c and 4d), the transmission of port 1(2) was measured as -0.7 *±* 0.8 dB (−11.8 *±* 0.8 dB) and -25.9 *±* 3.5 dB (−3.7 *±* 2.1 dB) for 473 nm and 638 nm light, respectively. On average, for TM-polarized light, the extinction ratio at 473 nm was 11.1 dB, while the extinction ratio at 638 nm was 22.2 dB.

**Fig 4.**
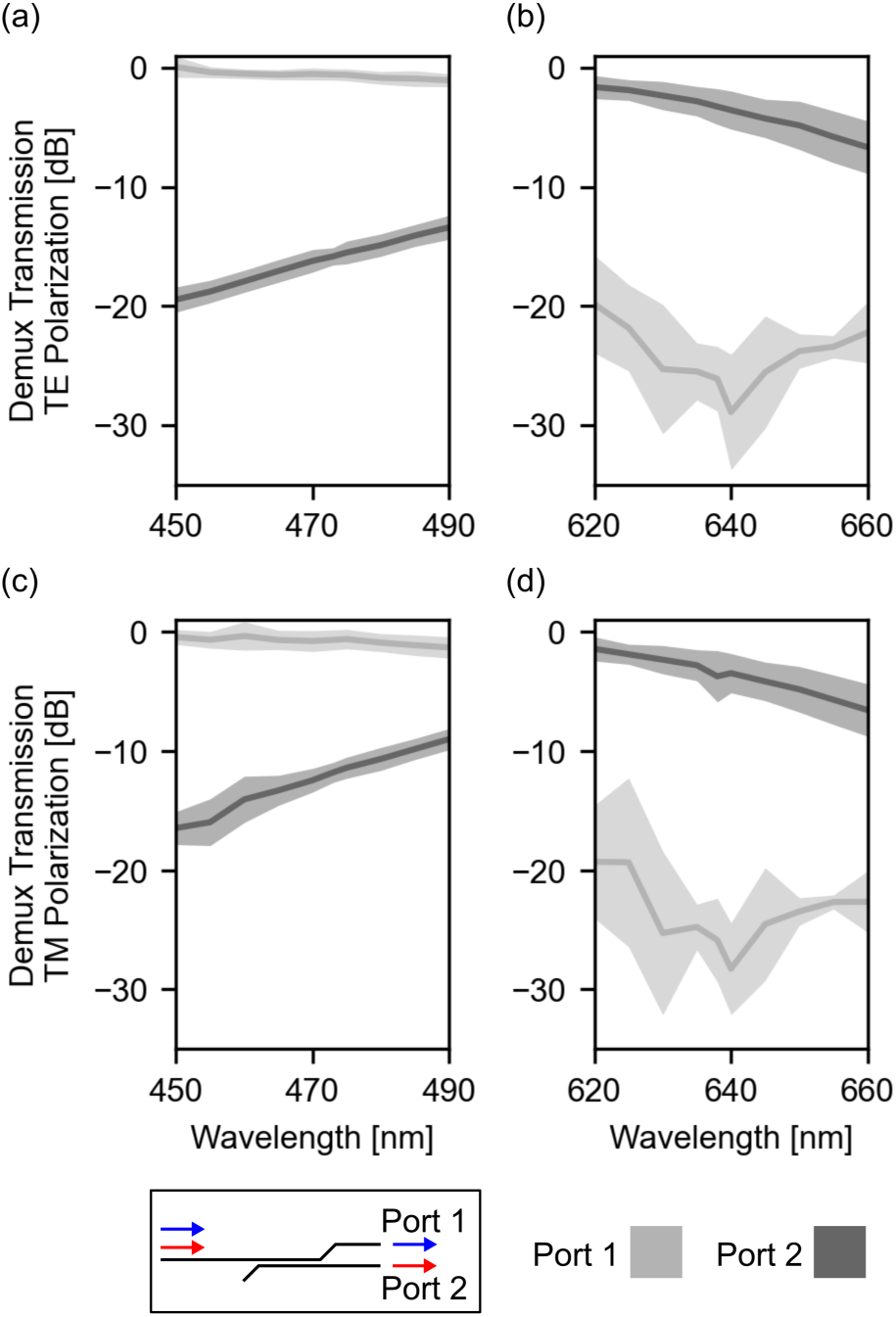
Characterization of wavelength demultiplexer test devices (n=5). For all panels, Port 1 was designed to selectively transmit 473 nm light and Port 2 was designed to selectively transmit 638 nm light. (a,b) Transmission measurements for TE-polarized light over wavelength spans of (a) 450 - 490 nm and (b) 620 - 660 nm. (c,d) Transmission measurements for TM-polarized light for wavelengths (c) 450 - 490 nm and (d) 620 - 660 nm.

To assess the region of excitation achieved by the blue and red grating coupler emitters, the grating coupler beam profiles were measured in fluorescent solution (Methods 4.6) and the in-plane and side profiles of the beams were evaluated. Representative micrographs of the in-plane/top profiles of the 473 nm and 638 nm beams are shown in Fig. 5a (top row), while representative micrographs of the side profiles of the 473 nm and 638 nm beams are shown in Fig. 5a (bottom row). Both the 473 nm and 638 nm beam profiles achieve a higher fill factor in the side profile by emitting two distinct lobes, emitted from the separate grating structures in SiN2 and SiN1 (Fig. S1). The full width at half maximum (FWHM) beam widths were measured at a distance 100 *µ*m away from the grating for the top and side profiles (n=7). The beam widths of the top profile were measured as 61 *±* 1 *µ*m and 91 *±* 2 *µ*m for the 473 nm and 638 nm beams, respectively (Fig. 5b). Similarly, the measured beam widths of the side profiles were 65 *±* 4 *µ*m and 54 *±* 1 *µ*m for the 473 nm and 638 nm beams, respectively (Fig. 5c).

**Fig 5.**
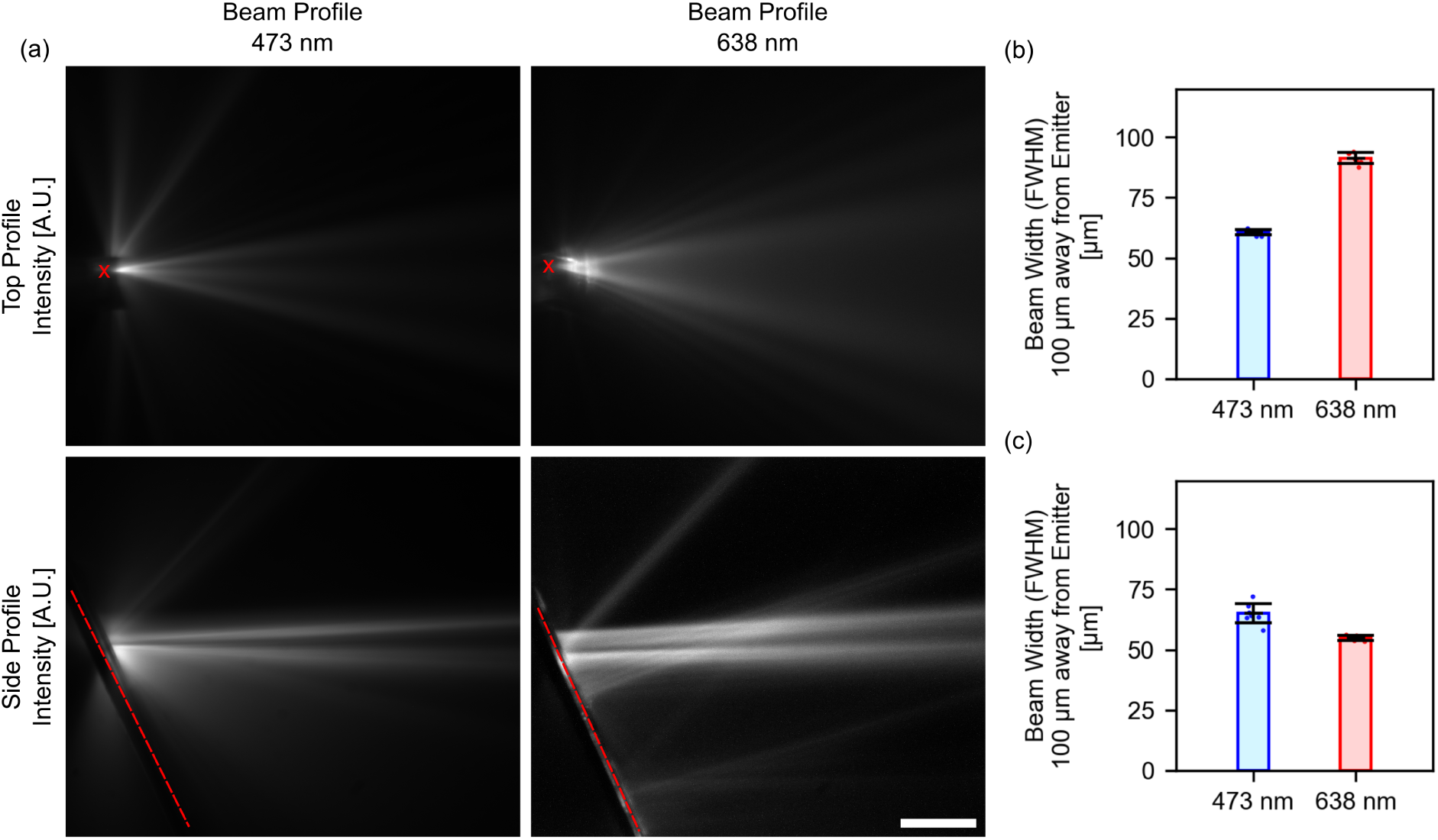
Beam profile measurements for 473 nm and 638 nm grating coupler emitters. (a) Representative normalized images showing the beam profiles for 473 nm light in fluorescein solution (left column) and 638 nm light in AlexaFluor647 solution (right column). The in-plane/top profile of the beams are shown in the top row and the side profile of the beams are shown in the bottom row. The neural probe shank is marked with a red x (red dashed line) in the top profile (side profile). Scale bar: 100 *µ*m. (b) Full width at half maximum (FWHM) beam width measurements for the 473 nm and 638 nm beam top profiles measured at 100 *µ*m away from the grating coupler (n=7). (c) FWHM beam width measurements for the 473 nm and 638 nm beam side profiles measured at 100 *µ*m away from the grating coupler (n=7).

### 2.3 In vivo evaluation with dual-color photostimulation and electrophysiological recording

To evaluate the performance of the neural probes, we conducted experiments in optogenetic mice responsive to either blue or red photostimulation. A nanophotonic neural probe was connected to a custom-designed dual-color laser scanning system (Fig. S2) via a 16-core multicore optical fiber^29^ (see Methods) for delivering programmatically-defined high-intensity blue and red light pulse trains (10 pulse trains, 10 pulses per pulse train, 30 ms pulse width, 5 Hz) through up to 16 of the available 26 emitter pairs. The nanophotonic probe was implanted along with a Neuropixels 2.0 single-shank probe^24^ to capture high-density electrophysiological data from the target brain region during dual-color photostimulation at multiple depths.

We assessed the functionality of blue photostimulation in cortex of a blue-light sensitive VGAT-ChR2 mouse (Fig. 6), which expresses Channelrhodopsin-2 in all GABAergic (inhibitory) interneurons. To maximize the blue photostimulation intensity, the end of the multicore fiber was moved into the blue laser focal position in the dual-color laser scanning system (Fig. S2d). At this position, the total system insertion loss was measured as 34.0 *±* 1.0 dB (41.7 *±* 0.7 dB) at 473 (638) nm, for an average maximum emitter power output of 119.4 *µ*W and 12.2 *µ*W for 473 nm and 638 nm light, respectively. Blue-light pulse trains were delivered to the cortex through emitter 1 (deep) to 5 (superficial) while electrophysiological data from the nanophotonic probe was monitored. Red photostimulation was also delivered as a control for the blue photostimulation. Electrophysiological data from the nanophotonic neural probe underwent spike sorting (Methods 4.9), however no high-quality units were detected. In addition to the low-density electrophysiological recordings captured by the nanophotonic probe, high-density electrophysiological recordings were captured using a Neuropixels probe placed nearby the nanophotonic probe. Representative data showing the average firing rate of all high-quality units (n=113) detected by the Neuropixels probe in response to photostimulation from emitter 3 are shown in Figs. 6a and 6b. During red photostimulation, no obvious effects on the firing rate were observed (Fig. 6a). In contrast, widespread reductions in firing rate were observed during blue photostimulation, consistent with synaptically driven inhibition of the local population by blue-light-sensitive GABAergic interneurons (Fig. 6b). A conceptual image showing the relative positions of the nanophotonic probe and Neuropixels probe is shown in Fig. 6c. Representative data showing the average firing rate of one unit in response to red and blue photostimulation from each emitter can be seen in Figs. 6d and 6e. Here, broad inhibition is seen for blue photostimulation from all emitters. Lastly, the average firing rate of all Neuropixels units was quantified for photostimulation through emitters 1 to 5 for comparison (Fig. 6f). For nearly all emitters, the average firing rate during blue photostimulation was significantly reduced compared to the firing rate during red photostimulation (emitters 1, 2, and 4: p*<*0.001, emitter 3: p=0.003, emitter 5: p=0.088, Mann-Whitney U rank test). Following the experiments, we attempted to modify the photostimulation pulse train settings for higher frequency (10 Hz pulse frequency and 10 ms pulse width). However, for the blue laser source, these settings resulted in irreparable damage to the selected emitters (1 and 2), likely due to light-induced damage to the UV-curing epoxy which bonded the multicore fiber to the probe.

**Fig 6.**
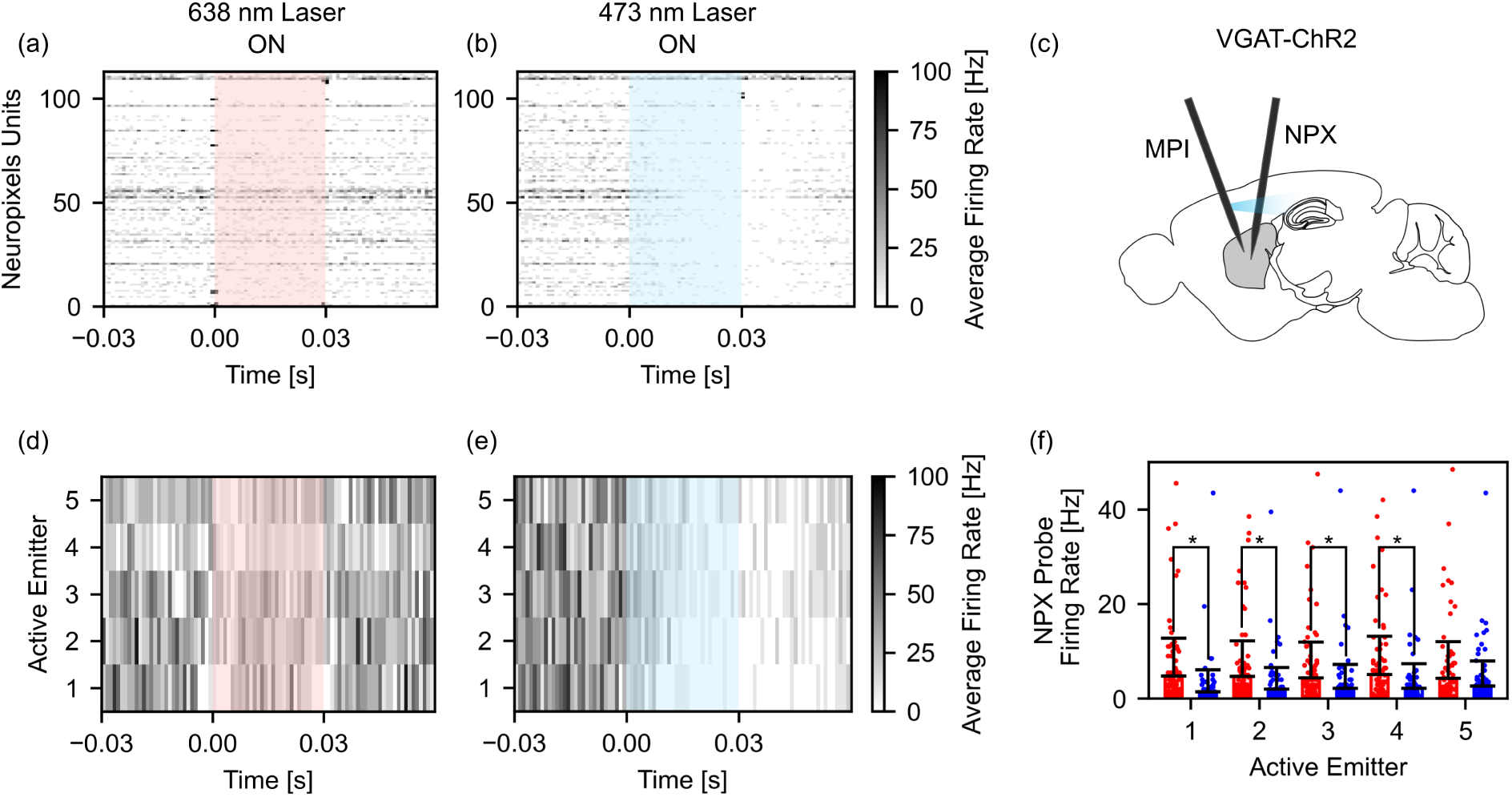
*In vivo* photostimulation experiment results in cortex of a VGAT-ChR2 mouse. (a,b) Representative spike-sorted data of Neuropixels units (n=113) during (a) red and (b) blue photostimulation. (c) Conceptual image showing the relative position of the nanophotonic (MPI) and Neuropixels (NPX) probes during the experiment. (d,e) Representative data from one Neuropixels unit during (d) red and (e) blue photostimulation from emitters 1 through 5. (f) Average firing rate data for Neuropixels units for photostimulation from emitters 1 through 5. The colors of the bars in (f) correspond to the color of photostimulation (left bar: red; right bar: blue).

Next, we assessed the functionality of red photostimulation in a mouse that expresses red-light-sensitive ChrimsonR in indirect medium spiny neurons of the striatum (Adora2a-Cre mouse injected with AAV-flex-ChrimsonR). To maximize the red photostimulation intensity, the end of the multicore fiber was moved into the red laser focal position in the dual-color laser scanning system (Fig. S2d). At this position, the total system insertion loss was measured as 43.3 *±* 0.5 dB (38.3 *±* 0.7 dB) at 473 (638) nm, for an average maximum emitter power output of 14.0 *µ*W and 26.6 *µ*W for 473 nm and 638 nm light, respectively. Red photostimulation pulse trains were delivered to the dorsal striatum and overlying cortex through emitters 3 (deep) to 12 (superficial) while evoked electrophysiological activity was measured by the nanophotonic probe and a nearby Neuropixels probe. Blue photostimulation was delivered as a control, with an understanding that high-intensity blue-light may still activate ChrimsonR+ neurons but with a lower yield. Representative images of the preprocessed electrophysiological data evoked during red and blue photostimulation from emitter 10 are shown in Figs. 7a and 7b. A conceptual image showing the relative positions of the nanophotonic probe and Neuropixels probe is shown in Fig. 7c. For red photostimulation, large amplitude spikes were observed on the recording electrodes nearest to the active emitter. This localized activity was not observed during blue photostimulation. Representative average firing rate data of spike-sorted units during photostimulation from emitter 10 can be seen in Figs. 7d and 7e. The approximate level of the active emitter is represented using a red or blue arrow. As shown in the preprocessed electrophysiological data, localized activity was observed during red photo-stimulation (Fig. 7d) but not blue photostimulation (Fig. 7e). We quantified this local activation by calculating the average firing rate of all units (n=11) as a function of the unit distance from the active emitter for photostimulation from emitters 3 to 12 (Fig. 7f). Units nearest to the active emitter had significantly higher firing rates for red photostimulation when compared to blue photo-stimulation (p=0.047, Mann-Whitney U rank test). Firing rate data from one representative unit in response to photostimulation from emitters 3 - 12 is shown in Figs. 7g and 7h. Again, we observed highly localized activation near one emitter for red photostimulation but not blue photostimulation. In addition to the nanophotonic probe neural recordings, electrophysiological data was measured further away from the neural probe using a Neuropixels 2.0 probe. Representative average firing rate data for the Neuropixels units (n=176) during red and blue photostimulation from emitter 10 can be seen in Figs. 7i and 7j. No clear differences were observed in the firing rate of any units during red and blue photostimulation. The average firing rate for all units was calculated during photostimulation from emitters 3, 5, 7, 9, and 11 (representing a photostimulation span of 1.5 mm) to quantify the regional effects of photostimulation (Fig. 7k). Overall, no differences in the average firing rate of units during red and blue photostimulation were observed. In comparison to the broad inhibition seen on the Neuropixels probe in the VGAT-ChR2 mouse, the lack of any widespread response to red light in this experiment could be due to the lower maximum light power from the red vs. blue emitters, or the more limited spread of the virus used to express ChrimsonR.

**Fig 7.**
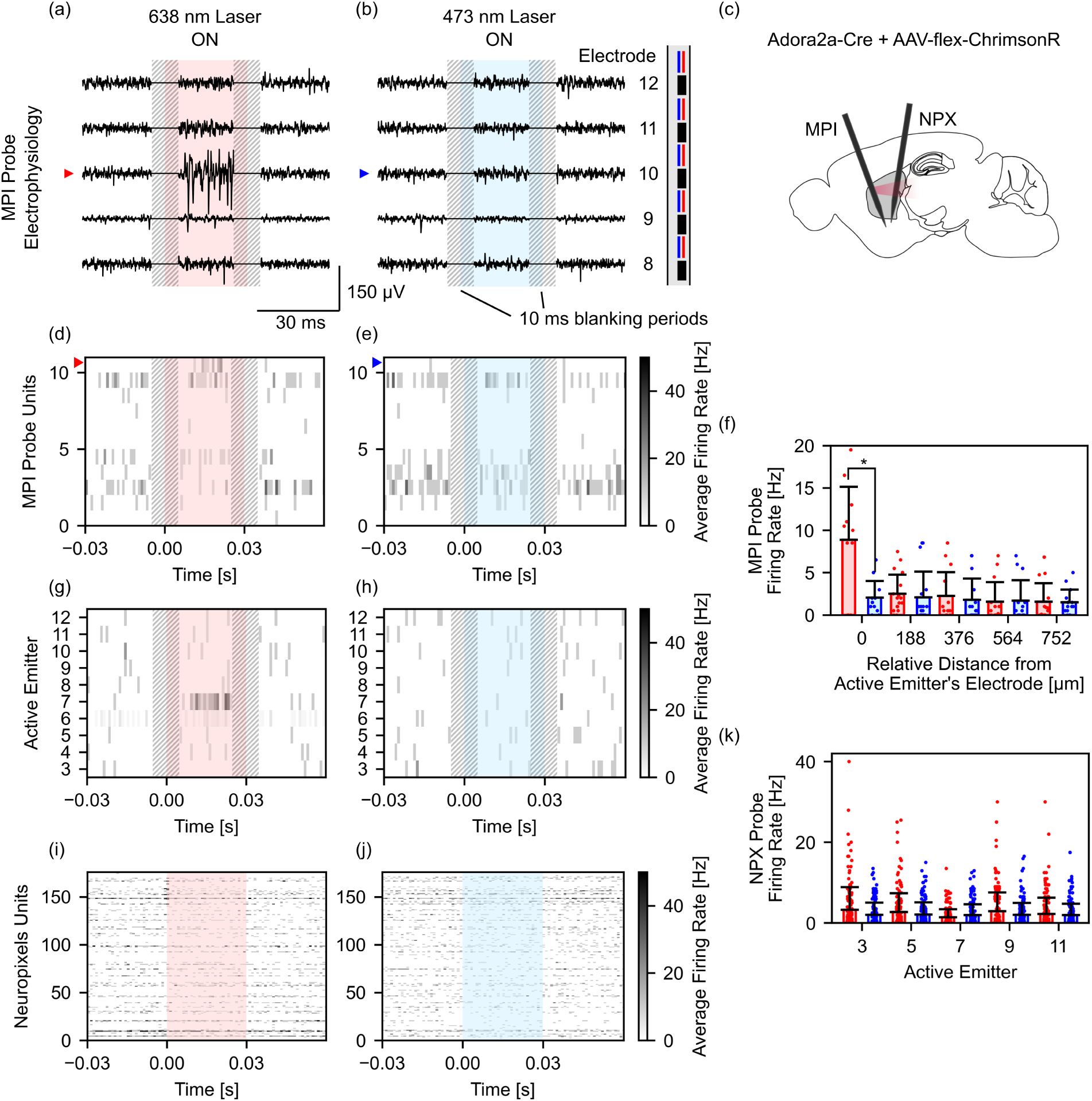
*In vivo* photostimulation experiment results in striatum of an Adora2a-Cre + AAV-flex-ChrimsonR mouse. (a,b) Representative preprocessed electrophysiological data from electrodes 8 to 12 on the nanophotonic probe during (a) red and (b) blue photostimulation. A 10 ms blanking period, centered on each pulse onset and offset, was applied for each stimulation pulse to remove stimulation artifacts. (c) Conceptual image showing the relative position of the nanophotonic (MPI) and Neuropixels (NPX) probes during the experiment. (d,e) Representative spike-sorted data of nanophotonic probe units (n=11) during (d) red and (e) blue photostimulation for one active emitter. (f) Average binned firing rate data for nanophotonic probe units as a function of the distance from the active emitter. (g,h) Representative firing data from one nanophotonic probe unit during (g) red and (h) blue photostimulation from emitters 3 through 12. (i,j) Representative spike-sorted data of Neuropixels units (n=176) during (i) red and (j) blue photostimulation. (k) Average firing rate data for Neuropixels units for photostimulation from emitters 3, 5, 7, 9, and 11. The color of the bars in (f) and (k) correspond to the photostimulation color (left bar: red; right bar: blue).

## 3 Discussion and Conclusion

We have characterized and demonstrated the functionality of a nanophotonic probe design with combined spatial multiplexing and on-shank wavelength-demultiplexers for high-density dual-color photostimulation and electrophysiological recording in two separate experimental settings: in a blue-light sensitive VGAT-ChR2 mouse and in a red-light sensitive Adora2a-Cre + AAV-flex-ChrimsonR mouse. In the current configuration, we connected 16 of the total 26 emitter pairs to a laser scanning system via a custom 16-core multicore fiber for red and blue photostimulation across a span of 2.92 mm - sufficient for stimulating and recording from cortex as well as deep brain structures in the mouse. Even in this limited configuration, this represents, to the authors’ knowledge, the most addressable emitters on a single photonic neural probe shank to date. A summary of our design in comparison to other state-of-the-art photonic and optoelectronic probe designs is shown in Table 1. With the combination of two multicore fibers, all 26 emitter pairs could be controlled, and the full 4.80 mm emitter span could be realized, which would improve this metric even further. We envision such high-density and long-span neural probes supporting optogenetic studies involving deep neural structures or neural systems spanning several millimeters, such as in cortex and deep brain structures like the basal ganglia,^30, 31^ hippocampus,^32^ or spinal cord.^33^ Because our nanophotonic neural probes are fabricated in a commercial Si photonics foundry with minimal post-processing, a direct path exists for scaling fabrication volumes to broadly disseminate these devices within the neuroscience community.

**Table 1.** Comparison of state-of-the-art photonic and optoelectronic neural probes for multicolor photostimulation and electrophysiological recording.

| Referenced Work | Ref. 19 | Ref. 17 | Ref. 12 | Ref. 13 | This work |
| --- | --- | --- | --- | --- | --- |
| Architecture | Photonic (SiN) | Photonic (SiN) | Optoelectronic (μLED) | Optoelectronic (OLED) | Photonic (SiN) |
| Emitter power | 39 μW (blue)<br>372 μW (red) | 5-10 μW (blue, red) | ∼200nW (blue, red) | 100 nW (blue)<br>40 nW (orange) | 212 μW (blue)<br>88 μW (red) |
| Total emitters | 14 x 2 colors | 3 x 2 colors | 16 x 2 colors | 1024 (orange or blue) | 26 x 2 colors |
| Emitters/shank | 14 x 2 colors | 3 x 2 colors | 16 x 2 colors | 256 (orange or blue) | 26 x 2 colors |
| Total electrodes | 960 (384 selectable) | 64 | 17 | 32 (bonded multielectrode array) | 26 |
| Electrodes/shank | 960 (384 selectable) | 64 | 17 | 32 (bonded multielectrode array) | 26 |
| Shank dimensions | 10 mm x 70 μm x 33 μm | 1 mm x 45 μm x 15 μm | 5 mm x 200 μm x 50 μm | 6.17 mm x 100 μm x 55 μm | 6.10 mm x 70 μm x 37 μm |
| Wavelengths | 450 nm, 638 nm | 450 nm, 655 nm<br>(central wavelengths) | 462 nm, 625 nm | 500 nm, 616 nm<br>(central wavelengths) | 473 nm, 638 nm |

We have shown that evanescent directional-coupler wavelength-demultiplexers provide an effective and compact method for demultiplexing two spectrally-distinct wavelengths of light (i.e., red and blue) to separate grating emitters on the neural probe shank. With an overall width of approximately 3 *µ*m, these devices are considerably more compact than microring resonator demultiplexers, which have diameters from 8 - 14 *µ*m^17^ (limited by waveguide bend losses). This compactness allows for more shank area to be allocated for waveguide routing. Moreover, the relatively broad bandwidth of directional-coupler devices reduces sensitivity to fabrication variations. For the directional-coupler wavelength-demultiplexer presented here, the average extinction ratio between output ports ranged from 11.1 dB to 22.8 dB, depending on the wavelength and polarization. In the case of lowest extinction ratio (i.e., for 473 nm TM-polarized light), approximately 10% of the input light may transmit to the red-light optimized grating coupler emitter instead of the blue-light optimized emitter, resulting in a second, lower-intensity beam being emitted by the emitter pair. Due to the close proximity of the red and blue emitters within each emitter pair, photostimulation would still be highly localized to the same stimulation depth, however the volume of photostimulation would be altered by the combined spatial profile of the two beams. In particular, the red-light emitter would emit 473 nm TM-polarized light at an angle of 35-40*^◦^*, broadening the side profile of the beam and resulting in a larger photostimulation volume (data not shown). To avoid this effect, multiple directional-coupler wavelength-demultiplexers could be cascaded before the emitter pair to improve the extinction ratio further and reduce emitter crosstalk.

We have demonstrated that high-intensity red and blue photostimulation pulse trains can be delivered with millisecond precision and average output powers of 119.4 *µ*W and 29.5 *µ*W for blue and red light, respectively, for a particular nanophotonic probe sample *in vivo*. Based on the average transmission of probes measured in this study (see Fig. 3a) and the maximum red and blue optical power available from the dual color scanning system (see Fig. S2d), the average maximum output power which could be achieved by the current design is 212 *µ*W for 473 nm light and 88 *µ*W for 638 nm light. These emitter powers are primarily limited by probe transmission, which itself is limited by variability in fiber to chip coupling. After packaging, probe transmission decreases by as much as 15 dB due to variable fiber to chip coupling, typically related to drift and misalignment of the multicore fiber relative to the on-chip edge couplers during or shortly after UV-curing. To reach the high output powers demonstrated in this study, we compensate for low probe transmission using high-power externally-coupled lasers, with output powers ranging from 180 mW - 300 mW. However, high-intensity laser light may also damage the packaged device. When adjusting the pulse train settings, increasing the frequency of the blue photostimulation pulse train resulted in emitter damage, likely due to damage to the UV-curing epoxy at the base of the nanophotonic probe. At a pulse frequency of 5 Hz, we did not observe any emitter damage during experiments. Compared to other technologies, this functional limitation is not unique, as *µ*LED optoelectrodes must also carefully control their pulse frequency and duty cycle when stimulating at high optical power to avoid thermal effects in neural tissue.^34, 35^ With optimization of the probe transmission, either through improved fiber to chip coupling or more efficient photonic structures, a lower laser power could be utilized to deliver similar output powers, which may reduce the chances of emitter failure during stimulation. Otherwise, alternative epoxies could be investigated with the goal of improved tolerance to high-power blue laser light.

The current probe design is able to record neural data with 26 electrodes across a wide span but with low resolution. In practice, these recordings can offer some insight into the local effects of photostimulation. However, when paired with high-density electrophysiology probes (as demonstrated with Neuropixels probes in the current study), the regional effects of stimulation can also be explored at greater resolution. In combination, tools separately optimized for either high-density photostimulation or high-density electrophysiological recording may offer researchers the ability to probe neural function at previously unachievable resolution and scale.

In future designs, we envision the emitter density of the nanophotonic neural probes improving further by employing a combination of denser waveguide routing, more compact demultiplexing structures, and multiple photonic layers for independent routing. Moreover, by using multicore fibers with more than 16 cores, a greater number of addressable photonic channels could be realized. Additionally, to transition from acute to chronic experimental configurations, further research is necessary to produce a reliable connector for the multicore fiber connected to the neural probe. In the current configuration, the multicore fiber remains permanently attached to the probe after packaging, limiting compatibility with chronic experimental designs, where the mouse would ideally be disconnected from the external laser scanning system outside of experiments. A reliable multicore fiber connector may permit the system to be disconnected/reconnected, permitting chronic probe implantation. Alternatively, on-chip Mach-Zehnder switching architectures^16, 19^ could be used to increase the number of addressable emitters while simplifying the fiber to chip connection.

In conclusion, we have demonstrated a novel, foundry-fabricated, nanophotonic neural probe capable of dual-color photostimulation and electrophysiological recording. Through a combination of spatial and wavelength multiplexing, notably using on-shank robust directional-coupler wavelength-demultiplexers and low-crosstalk waveguide arrays with dissimilar widths, a record 52 grating couplers (26 blue and 26 red) were integrated onto a single narrow (70 *µ*m wide, 37 *µ*m thick) neural probe shank, which also included 26 recording electrodes. Paths toward further increasing emitter density and improving optical transmission have been detailed. Our *in vivo* experiments in mice expressing red- and blue-light sensitive opsins have demonstrated the functionalities of the neural probe, while also highlighting the practicality of multi-probe experiments using photonic neural probes for photostimulation and CMOS-based probes for electrophysiological recording, each independently designed for either high photonic emitter or electrode density. Overall, this work provides a foundation for further scaling and maturation of dual-color photonic neural probes - toward higher emitter densities, larger optical output powers, and volume fabrication.

## 4 Methods

### 4.1 Neural probe fabrication

Neural probes were fabricated on 200-mm diameter Si wafers at Advanced Micro Foundry (AMF) using a two-layer SiN visible-light integrated photonics platform with 3 Al metal layers.^25^ Fabrication began with deposition of the silicon dioxide (SiO_2_) waveguide bottom cladding. Next, 120-nm (SiN1) and 75-nm (SiN2) thick SiN waveguides and their interlayer SiO_2_ layer were defined by plasma enhanced chemical vapor deposition, 193 nm deep ultraviolet photolithography, and reactive ion etching; chemical mechanical polishing was used for planarization. Next, the SiO_2_ top cladding of the waveguides, three Al metal layers (M1, M2, M3) with interconnecting vias were formed. Recording electrodes were coated with titanium nitride (TiN). Deep trenches were etched to define the probe shapes and edge coupler facets. Finally, backgrinding was performed to thin the wafers to *∼*100 *µ*m and separate the chips.

### 4.2 Neural probe post-processing and laser-induced electrode impedance reduction

Neural probes were mechanically polished to reduce the probe shank thickness from approximately 100 *µ*m to 40 *µ*m (NOVA Optical Polishing System, Krell Technologies Inc., Neptune City, NJ, USA). Samples were encased in mounting wax to avoid device damage during polishing. Samples were lowered at a rate of 5 *µ*m every 30 seconds against a rotating 9 *µ*m grit-size polishing film until a final thickness was reached. The thickness of neural probe samples was measured optically by imaging sample side profiles under a microscope with respect to a 1 mm measurement scale. All measurements were compared using Fiji.^36^ Following polishing, samples underwent at least two 16-hour acetone bath treatments to reduce surface contamination from the mounting wax.

Laser-induced electrode impedance reduction was achieved using previously described methods.^18^ Briefly, a 2-photon microscope (Bruker Corporation, Billerica, MA, USA) equipped with a femtosecond laser (Monaco, Coherent Corporation, Saxonburg, PA, USA) was used to roughen electrode surfaces via selective laser scanning. The femtosecond laser repetition rate was set to 10 kHz, the wavelength was set to 1035 nm, and the average incident power was set to 60-80 *µ*W. A laser scanning sequence was applied once per electrode, and repeated until the electrode surface appeared visibly darkened when viewed under an optical microscope (see Fig. 3c, right).

### 4.3 Neural probe packaging

Flexible printed-circuit boards (PCBs) were designed and fabricated (Würth Elektronik, Niedernhall, Germany) for connecting the neural probes to an external data acquisition system. Custom aluminum metal probe holders were designed and fabricated (Proto Labs Germany GmbH, Putzbrunn, Germany) for mounting the neural probes and flexible PCBs. Custom aluminum metal stereotactic adapters were designed and fabricated in house by the Allen Institute for implanting the neural probes. Neural probe samples were mounted on the aluminum probe holder assemblies using metal epoxy (Loctite Ablestik 84-1LMIT1, Henkel AG & Co. KGaA, Düsseldorf, Germany), which was cured at 130 *^◦^*C for 4 hours. Neural probe electrodes were then connected to the PCB via Al-wire wirebonding, which were encapsulated in dielectric epoxy (Katiobond GE680, Delo, Germany). After electrical packaging, neural probe samples were connected to a custom 16-core multicore fiber (Corning Inc., Corning, NY, USA) using previously described methods.^18^ Briefly, multicore fibers were aligned to the neural probe edge couplers (each core coupled to an on-chip edge coupler) using a 6-axis alignment stage and bonded using UV-curable optical epoxies (OP-67-LS and OP-4-20632, Dymax Corp., Torrington, CT, USA). After curing, the optical epoxy was coated in an optically opaque epoxy (EPO-TEK 320, Epoxy Technology, Billerica, MA, USA) to reduce stray light emission.

### 4.4 Electrode impedance measurements

Electrode impedance values for neural probe chips were measured using an impedance analyzer (Keysight E4990A, Keysight Technologies, Santa Rosa, CA, USA). Electrodes were lowered into a bath of 1x Dulbecco’s phosphate buffered saline (DPBS) (Merck KGaA, Darmstadt, Germany) and electrode bond pads were connected to the impedance analyzer using a tungsten needle. The electrode impedance was measured using a 10 mV waveform relative to a Ag/AgCl electrode. After packaging, electrode impedances were measured using the impedance measurement function of an Intan RHD2132 headstage (Intan Technologies, Los Angeles, CA, USA). Packaged neural probes were immersed in 1x DPBS and electrode impedances were measured relative to a Ag/AgCl electrode.

### 4.5 Wavelength demultiplexer measurements

The optical transmission of wavelength demultiplexer test devices (on separate test chips) were measured using a supercontinuum laser (SuperK Extreme, NKT Photonics, Birkerød, Denmark) and an optical power meter (Newport, Irvine, CA, USA). The supercontinuum laser output was po-larized using a polarization filter and the output polarization was maintained using a polarization-maintaining (PM) fiber. The output wavelength of the supercontinuum laser was set using a tunable optical bandpass filter (LLTF Contrast, NKT Photonics). A separate length-matched waveguide test structure was measured to subtract the loss of the edge couplers and routing waveguide from the demultiplexer test structures.

### 4.6 Beam profile and transmission measurements

Neural probe optical beam profiles were measured in fluorescent solution to characterize the in-plane and side profile of the beam. The top profiles were measured using a vertically-positioned fluorescence microscope (Objective: 10x Plan Apo, Mitutoyo Corporation, Kawasaki, Japan). Side beam profiles were measured using an additional horizontally-positioned fluorescence microscope (Objective: 5x Plan Apo, Mitutoyo). Packaged neural probe samples were connected to a custom laser scanning system with a depolarized supercontinuum laser input (SuperK Fianium, NKT Photonics). The emission wavelength of the supercontinuum laser was selected using a tunable optical bandpass filter (LLTF Contrast, NKT Photonics). The laser scanning system used for optical characterization of the probes was different from the dual-color scanning system used for *in vivo* experiments in this work (Sec. 4.7); having a single-mode fiber input and no free-space depolarizer. This simpler scanning system is detailed in our previous work.^37^ To characterize the blue grating coupler emitter, the optical input was set to 473 nm and the probe was immersed in 100 *µ*M fluorescein solution. Optical emission filters (ET525/50M, Chroma Technology Corp., Bellows Falls, VT, USA; BrightLine Basic Fluorescence Filter 525/39, IDEX Corporation, Northbrook, IL, USA) were inserted in the vertically-positioned and horizontally-positioned fluorescence microscopes, respectively, and the neural probe beam profiles were imaged (via imaging the fluorescence excited by the neural probe output). To characterize the red grating coupler emitter, the optical input was set to 638 nm and the probe was immersed in 2 mg/L AlexaFluor647 solution (ThermoFisher Scientific, Waltham, MA, USA). Optical emission filters (FELH0650 - 650 nm longpass filter, Thorlabs, Inc., Newton, NJ, USA) were inserted in the vertically-positioned and horizontally-positioned fluorescence microscopes for fluorescence imaging of the beam profiles. Probe transmission for three packaged nanophotonic probes were measured using the supercontinuum laser source and an optical power meter (Newport, Irvine, CA, USA). An additional packaged nanophotonic probe, which was used in *in vivo* experiments, used the custom dual-color laser scanning system during transmission measurements.

### 4.7 Custom dual-color laser scanning system

A custom free-space laser scanning system (Fig. S2a and S2b), similar to that described in Ref. 18, was designed for delivering high-intensity blue- and red-light to a multicore fiber coupled to the neural probe. A 300 mW 473 nm laser diode (Cobolt 06-MLD, Hübner Group, Kassel, Germany) and 180 mW 638 nm laser diode (Cobolt 06-MLD, Hübner Group) were directed through separately-controlled variable neutral density filters (NDC-25C-2-A, Thorlabs, Inc., Newton, NJ, USA) and achromatic have-wave plates (AHWP10M-600, Thorlabs, Inc.) in rotation mounts. The beams were then combined using a dichroic mirror and the combined beam was sent through a free-space Mach-Zehnder interferometer depolarizer with an optical path length difference of approximately 220 mm. The depolarizer minimized the effects of polarization variability due to temperature and stress fluctuations in the multicore fiber, which additionally varies between cores (each core has a separate Jones matrix). Since the neural probes function for both TE- and TM-polarized light (albeit with higher performance for the TE-polarization), the probes also function with depolarized light, which is effectively equal parts TE- and TM-polarized on-chip. After depolarization, the beam was reflected off a right-angle prism mirror and directed onto a MEMS mirror (Mirrorcle Technologies Inc., Richmond, CA, USA) followed by custom focusing optics. The focused beam was coupled into the facet of the multicore fiber, which was positioned using a 3-axis alignment stage. The MEMS mirror was controlled by a custom MATLAB GUI (MathWorks, Natick, MA, USA) to direct the beam at one of the 16 cores of the multicore fiber (i.e., the position of the MEMS mirror selected the fiber core and corresponding on-shank grating coupler pair to emit light). Pulse trains were defined via modulation of the red and blue lasers. The custom scanning optical system was originally designed for single-color operation, so in the current version there is a chromatic focus shift between the red and blue laser wavelengths. Thus, for experiments, the axial position of the multicore fiber was positioned at either the blue-beam focus, red-beam focus, or an intermediate balanced defocus position between the blue- and red-beam foci depending on whether greater blue or red photostimulation intensities were required. Full chromatic correction can be achieved in the future by adding a diffractive optical element to the existing custom scanning optics design in order to maximize efficiency for red and blue light simultaneously.

### 4.8 Animal experiments

All animal experiments were conducted under protocols approved by the Institutional Animal Care and Use Committee (IACUC) at the Allen Institute in Seattle, WA. A VGAT-ChR2 mouse (male) and an Adora2a-Cre mouse (male) injected with AAV-flex-ChrimsonR were used for assessing the functionalities of a packaged neural probe *in vivo*. Under deep isoflurane anesthesia, a titanium headframe was attached to the skull with Metabond (Parkell, Inc.). A craniotomy and durotomy were performed over most of the left dorsal skull, which was replaced with a custom 3D printed cranial window.^38^ For the Adora2a-Cre mouse, 300 nL of virus (AAV-flex-ChrimsonR) was injected into dorsal striatum at a coordinate of 0.7 mm posterior, 2.45 mm lateral, and 2.6 mm ventral to the bregma skull suture. Following at least two weeks of recovery and at least 24 h prior to recording, a protective layer of durable silicone (SORTA-Clear, Smooth-On, Inc.) was replaced with a soft silicone elastomer (”Dura-Gel”, DOWSIL 3-4680) that would allow probes to penetrate into the brain.

Mice were trained to sit quietly while head-fixed in a 30 cm diameter plastic tube over the course of three habituation sessions. On the day of the recording, a nanophotonic probe and Neuropixels 2.0 single-shank probe were mounted on separate modules of a custom insertion system. Modules were placed at 4° from vertical along the anterior-posterior axis and either 0-5° (nanophotonic probe) or 25-32° (Neuropixels probe) along the medial-lateral axis. Each module included a 3-axis manipulator (M3-LS-3.4-15, New Scale Technologies) that allowed each probe to be moved independently with sub-micron precision. Probes were visualized using a pair of long-working-distance microscopes (InfiniProbe, Infinity Photo-Optical). Once the mouse was head-fixed, probes were aligned with holes in the cranial window so they would cross within 300 *µ*m of one another in the target structure (retrosplenial cortex in VGAT-ChR2 mouse and dorsal striatum in the Adora2a-Cre mouse). Probes were inserted into the brain at a rate of 200 *µ*m/min. Electrophysiological data from the Neuropixels probe was recorded using a Neuropixels basestation in a National Instruments PXI chassis. Electrophysiological data from the nanophotonic neural probes was digitized using a 32-channel recording headstage (RHD2132, Intan Technologies) connected to an Open Ephys acquisition board (Open Ephys Production Site, Lisbon). All data was visualized and recorded using the Open Ephys GUI.^39^

### 4.9 Electrophysiological data analysis

Electrophysiological data was losslessly compressed using the WavPack algorithm^40^ and uploaded to an Amazon S3 storage bucket. Neuropixels data was processed using the SpikeInterface package^41^ running in the Allen Institute for Neural Dynamics Code Ocean computing environment. Data was high-pass filtered at 300 Hz and phase-shifted to align samples multiplexed into the same analog-to-digital converter (ADC), and a common median reference (CMR) was applied across all channels. Data was sorted using Kilosort 2.5,^24^ followed by calculation of standard quality metrics.^31^ All units included in the analysis had *≥* 200 spikes, inter-spike interval ratios *<* 0.5 and median spike amplitudes *>* 50 *µ*V.

Electrophysiological data from the nanophotonic probe was analyzed using a custom electrophysiology processing pipeline in SpikeInterface^41^ at the Max Planck Institute of Microstructure Physics. Data was preprocessed by applying a band pass filter (300 Hz to 6000 Hz) and a common median reference. To remove photostimulation artifacts, a 10 ms blanking period, centered on each stimulation pulse onset and offset, was applied. Spike sorting was completed using SpyKING CIRCUS2, an internal spike sorter available in SpikeInterface based on the SpyKING CIRCUS spike sorter.^42^ After sorting, spike and quality metrics were calculated for each unit. Units were manually curated in Phy.^43^ Units with less than 200 spikes and units with noise-like waveforms were excluded from the analysis. All units with similarity scores *≥* 0.9 were compared and considered for manual merging. All units included in the analysis had inter-spike interval ratios *<* 0.5 and median spike amplitudes *>* 50 *µ*V.

### 4.10 Statistical analysis

Average binned firing rate data from the *in vivo* experiments during blue and red photostimulation were compared with Mann-Whitney U rank tests using the SciPy Python package. Significance is indicated for p*<*0.05. All reported values are mean *±* SD unless otherwise stated.

## Code and Data Availability

Electrophysiological data reported in this study and code for analyzing data are available from David A. Roszko upon reasonable request.

## Acknowledgments

This work was supported by the Max Planck Society. The authors thank Andrei Stalmashonak at the Max Planck Institute of Microstructure Physics for his advice and assistance with optical measurements.

**David A. Roszko** is pursuing a PhD at the University of Toronto (Toronto, Canada), with research conducted at the Max Planck Institute of Microstructure Physics (Halle (Saale), Germany). He received his BSc degree in electrical engineering and MSc degree in neuroscience from the University of Alberta (Edmonton, Canada) in 2018 and 2021, respectively. His current research interests include the design, fabrication, and characterization of nanophotonic neural interfaces for multicolor photostimulation.

**Fu-Der Chen** received his BASc (2017) and MASc (2019) degrees in Electrical and Computer Engineering from the University of Toronto, Ontario, Canada. He is pursuing a PhD degree in Electrical and Computer Engineering at the University of Toronto while conducting his research at the Max Planck Institute of Microstructure Physics and the Krembil Brain Institute. His research interest is to develop implantable silicon probes with integrated photonic devices for optogenetic stimulation and microelectrodes for electrophysiological recording.

**John Straguzzi** received his MSc in electrical and computer engineering from Drexel University, Philadelphia PA, USA in 2015. From 2015 to 2018, he worked as a research and development engineer for Agilent Technologies, Wilmington DE, USA. He has been a scientific research staff member at the Max Planck Institute of Microstructure Physics, Halle (Saale), Germany, since 2019, where he focuses on digital systems and PCB design to support the group’s experiments on photonic integrated circuits.

**Hannes Wahn** received his MSc degree in optical engineering from the University of Applied Sciences, Cologne, Germany, in 2015. From 2015 to 2020, he worked as an optical development engineer in industry. Since 2020, he is a research staff member at the Max Planck Institute of Microstructure Physics. His current research is focused on optical system design for applications such as microscopy, laser beam delivery and displays, as well as computational imaging microscopy for neuroscience applications.

**Alec Xu** is a PhD student at the University of Oxford, working on new liquid crystal devices for miniaturized adaptive optics, work and interests that were sparked during an internship at the Max Planck Institute of Microstructure Physics. Alec holds a BASc in Engineering Physics from the University of Toronto.

**Blaine McLaughlin** received his BSc degree in astrophysics and his MSc degree in physics from the University of Calgary, Alberta, Canada, in 2017 and 2022, respectively. Since 2024, he is a silicon photonics engineer at the Max Planck Institute of Microstructure Physics. His current research is focused on the design and testing of silicon nitride photonic devices for visible wavelengths.

**Xinxin Yin** is a research scientist at the Allen Institute. He earned his PhD in systems neuroscience from Tsinghua University and BS in biotechnology from the University of Electronic Science and Technology of China. His current research focuses on understanding the multi-regional dynamics of the flexible behavior at the scale of the entire brain.

**Hongyao Chua** received his bachelor degree in MSE from NTU, Singapore, in 2010. He joined IME, A*STAR in 2011, as a research engineer for silicon photonics technology development. He joined AMF as a senior research engineer in 2017, focusing on silicon photonics commercialization. He is currently a principal research engineer at AMF, working on silicon photonics manufacturing. He has authored/co-authored multiple peer-reviewed journal and conference papers.

**Xianshu Luo** received his PhD in ECE from HKUST, Hong Kong, in 2010. He joined IME, A*STAR in 2010, as a scientist for silicon photonics technology development. He joined AMF as co-founder and research director in 2017, focusing on silicon photonics commercialization. He is now in IME as a principal scientist working on advanced photonics. He has authored/co-authored more than 200 peer-reviewed journal and conferences papers and 5 book chapters and holds more than 20 patents.

**Guo-Qiang Lo** earned his PhD in electrical and computer engineering from the University of Texas (Austin, USA) in 1992. He has more than 30 years of experience in semiconductor manufacturing and optoelectronics in the USA and Singapore. Dr. Lo co-founded Advanced Micro Foundry in 2017 and now serves as CTO. He has received multiple prestigious awards, including the IEEE George E. Smith Award (2007) and Singapore’s National Technology and President Technology Awards (2008, 2010).

**Joshua H. Siegle** is a Senior Scientist at the Allen Institute for Neural Dynamics in Seattle. He received a PhD in Neuroscience from MIT in 2014. He now leads a group that is developing new techniques for recording and manipulating spiking activity across the mouse brain.

**Joyce K. S. Poon** is a Professor of Electrical and Computer Engineering at the University of Toronto and an Honorary Professor at the Technical University of Berlin. From 2018 to July 2024, she was a Director at the Max Planck Institute of Microstructure Physics. Since May 2024, she is the head of Photonics Architecture at Lightmatter Inc.. She specializes in integrated photonics on silicon. Prof. Poon is an Optica Fellow and a Fellow of the IEEE.

**Wesley D. Sacher** is an independent research group leader at the Max Planck Institute of Microstructure Physics. He received his PhD in electrical and computer engineering from the University of Toronto in 2015. From 2015 to 2018, he was a postdoctoral scholar at the California Institute of Technology. His current research is focused on integrated photonics for visible wavelengths and neurotechnologies for optogenetics and functional imaging.

---

**Fig S1.**
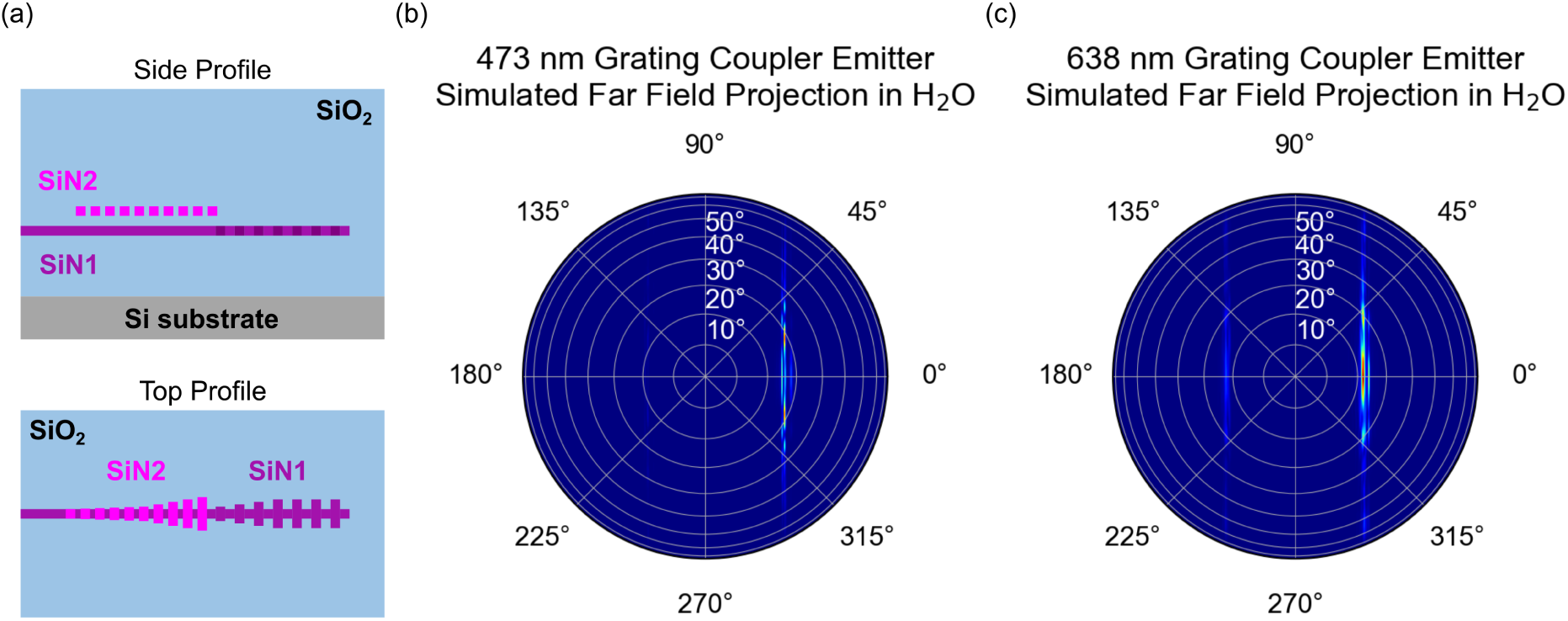
(a) Conceptual diagram of bi-layer grating coupler emitters. The design features a fully-etched grating coupler in SiN2, followed by a corrugated grating in SiN1. The 473 nm emitter (length: 101 *µ*m) consists of a SiN2 grating (fill factor: 50%, period: 0.47 *µ*m) with width apodization from 0.254 *µ*m to 2.038 *µ*m, followed by a SiN1 corrugated grating (fill factor: 50%, period: 0.47 *µ*m) with central waveguide width of 0.22 *µ*m and grating width apodization from 1.431 *µ*m to 1.5 *µ*m. The 638 nm emitter (length: 101 *µ*m) consists of a SiN2 grating (fill factor: 50%, period: 0.63 *µ*m) with width apodization from 0.493 *µ*m to 1.381 *µ*m, followed by a SiN1 corrugated grating (fill factor: 50%, period: 0.63 *µ*m) with central waveguide width of 0.22 *µ*m and grating width apodization from 0.627 *µ*m to 1.26 *µ*m. (b) Simulated far field projection of the 473 nm grating coupler emitter in water (refractive index, n = 1.3361). (c) Simulated far field projection of the 638 nm grating coupler emitter in water (refractive index, n = 1.3315). Simulations in (b,c) used Lumerical 3D FDTD with combined TE-and TM-polarized input light (modeling depolarized input light).

**Fig S2.**
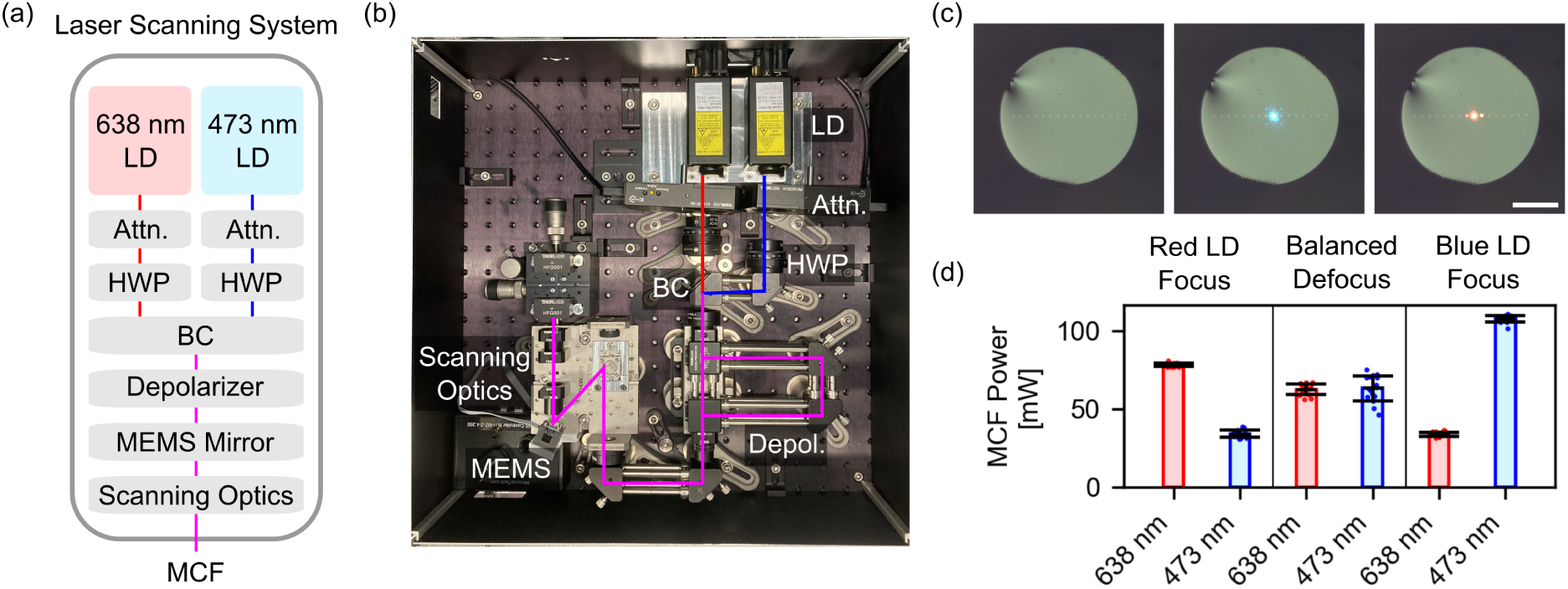
(a) Block diagram of custom dual-color laser scanning system. LD - laser diode, Attn. - variable attenuator, HWP - half-wave plate, BC - beam combiner, MEMS - microelectromechanical system, MCF - 16-core multicore fiber. (b) Labeled photograph of laser scanning system. (c) Micrographs of a 16-core multicore fiber facet without (left) and with blue/red (center/right) laser light coupled to a single core. Scale bar: 100 *µ*m. (d) Average available output power from MCF connected to the laser scanning system as a function of MCF axial position. The MCF can be aligned to either the red LD or blue LD focal points for maximum power at a single wavelength, or can be aligned to a balanced defocused position for approximately equal red and blue optical power.

